# Allopatric divergence, local adaptation, and multiple Quaternary refugia in a long-lived tree (*Quercus spinosa*) from subtropical China

**DOI:** 10.1101/112375

**Authors:** Li Feng, Yan-Ping Zhang, Xiao-Dan Chen, Jia Yang, Tao Zhou, Guo-Qing Bai, Jiao Yang, Zhong-Hu Li, Ching-I Peng, Gui-Fang Zhao

## Abstract

- The complex geography and climatic changes occurring in subtropical China during the Tertiary and Quaternary might have provided substantial opportunities for allopatric speciation. To gain further insight into these processes, we reconstruct the evolutionary history of *Quercus spinosa,* a common evergreen tree species mainly distributed in this area.
- Forty-six populations were genotyped using four chloroplast DNA regions and 12 nuclear microsatellite loci to assess genetic structure and diversity, which was supplemented by divergence time and diversification rate analyses, environmental factor analysis, and ecological niche modeling of the species distributions in the past and at present.
- The genetic data consistently identified two lineages: the western Eastern Himalaya-Hengduan Mountains lineage and the eastern Central-Eastern China lineage, mostly maintained by populations’ environmental adaptation. These lineages diverged through climate/orogeny-induced vicariance during the Neogene and remained separated thereafter. Genetic data strongly supported the multiple refugia (per se, interglacial refugia) or refugia within refugia hypotheses to explain *Q. spinosa* phylogeography in subtropical China.
- *Q. spinosa* population structure highlighted the importance of complex geography and climatic changes occurring in subtropical China during the Neogene in providing substantial opportunities for allopatric divergence.

## Introduction

Historical processes such as geographic and climatic changes have profoundly shaped the population genetic structure and demographic history of extant species (Hewitt, 2000, 2004). Climatic changes and heterogeneous environments could also provide opportunities for genetic divergence and diversification through adaptation to local or regional environments (Rainey & Travisano, 1998). When populations are adapted to dissimilar habitats, gene flow among them could be limited by selection and this might indirectly influence the whole genome, promoting neutral divergence through increased genetic drift (Wright, 1931; Nosil *et al.,* 2005).

Numerous studies have considered the effects of climatic changes since the Tertiary (e.g., Li *et al.,* 2013; Liu *et al.,* 2013; Wang *et al.,* 2015), but only a few have disentangled the roles of isolation by environment (IBE) and isolation by distance (IBD) in the population genetic divergence of temperate species (Mayol *et al.,* 2015; Zhang *et al.,* 2016). However, lack of exploring the roles of geographic and environmental forces in driving genetic structure and for inferring species’ past demography, may hinder the accurate and precise inference of the distinct roles of IBD and IBE in population genetic structure and in species’ demographic scenarios (Mayol *et al.,* 2015). Until recently, multiple matrix regression with randomization (MMRR) provided a robust framework, allowing powerful inferences of the different effects of IBD and IBE (Wang, 2013), and approximate Bayesian computation (ABC), allowing us to evaluate the most plausible demographic scenario and estimating the divergence and/or admixture time of the inferred demographic processes with a relatively low computation effort (Beaumont, 2010).

Wu & Wu (1998) suggested most of the Chinese flora could be divided into three subkingdoms (Sino-Japanese Forest, Sino-Himalayan Forest, and Qinghai-Xizang Plateau), all of are which included in subtropical China (21–34° N in South China). This region has a mild monsoon climate, complex topography, and high species diversity (Myers *et al.,* 2000; Qian & Ricklefs, 2000). Several hypotheses have been proposed to explain high species diversity here, among which the best known are those of Qian & Ricklefs (2000) and Harrison *et al.* (2001). Qian & Ricklefs (2000) suggested that the numerous episodes of evolutionary radiation of temperate forests, through allopatric divergence and speciation driven by mid-to-late Neogene and Quaternary environmental changes, promoted species diversity; species would have spread to lower elevations and formed a continuous band of vegetation during glacial periods and retreated to “interglacial refugia” at higher elevations during warmer periods. However, according to palaeovegetation reconstructions based on fossil and pollen data, Harrison *et al.* (2001) suggested that temperate forests were considerably less extensive than today and would have retreated southward to *ca.* 30° N during the Last Glacial Maximum (LGM), being replaced by non-forest biomes or by boreal and temperate-boreal forests.

In general, phylogeographic studies in subtropical China have reinforced the allopatric speciation hypothesis for species diversity, indicating a general pattern of multiple refugia and little admixture among refugial populations throughout glacial-interglacial cycles (Qiu *et al.,* 2011; Liu *et al.,* 2012). However, most of these studies focused on endangered species or temperate deciduous species with limited distribution, and only a few (Shi *et al.,* 2014; Xu *et al.,* 2014; Wang *et al.,* 2015) considered temperate evergreen species. Thus, further investigations are necessary to verify if this pattern of multiple refugia and allopatric speciation are applicable to the typical and dominant evergreen species inhabiting the temperate zone.

*Quercus spinosa* David ex Franch, belonging to the group *Ilex* (syn. *Quercus* subgenus *Heterobalanus*) within family Fagaceae, is a long-lived, slow-growing tree inhabiting East Asian temperate evergreen forests. This tree species could offer additional advantages to investigate the impacts of IBE and IBD compared to short-lived trees and buffer the effects of changes in population genetic structure due to its life history traits (e.g., longevity, overlapping generations, prolonged juvenile phase) (Austerlitz *et al.,* 2000). Its distribution range extends from eastern Himalaya to Taiwan, and is commonly found in subtropical China (Fig. 1a), growing on slopes and cliffs, in low-to mid-elevation (900–3800 m above sea level) (Wu *et al.,* 1999; Menitskii & Fedorov, 2005). Based on recent molecular phylogenetic evidence (Denk & Grimm, 2009, 2010; Hubert *et al.,* 2014; Simeone *et al.,* 2016), the Group *Ilex* (*c*. 30 spp.) is monophyletic and diversified rapidly during the late Oligocene/early Miocene, suggesting *Q. spinosa* originated during the Miocene. In addition, recent studies suggested a significant role for environmental adaptation in the origin and maintenance of genetic divergence among forest lineages (e.g., Mayol *et al.,* 2015; Ortego *et al.,* 2015; Sexton *et al.,* 2016) and Petit *et al.* (2013) suggested Fagaceae as ideal models for integrating ecology and evolution. Hence, this species provides an ideal model for investigating the intraspecific divergence and evolutionary dynamics of an evergreen forest species in subtropical China subjected to geologic and climatic changes since the Tertiary.

**Fig. 1.**
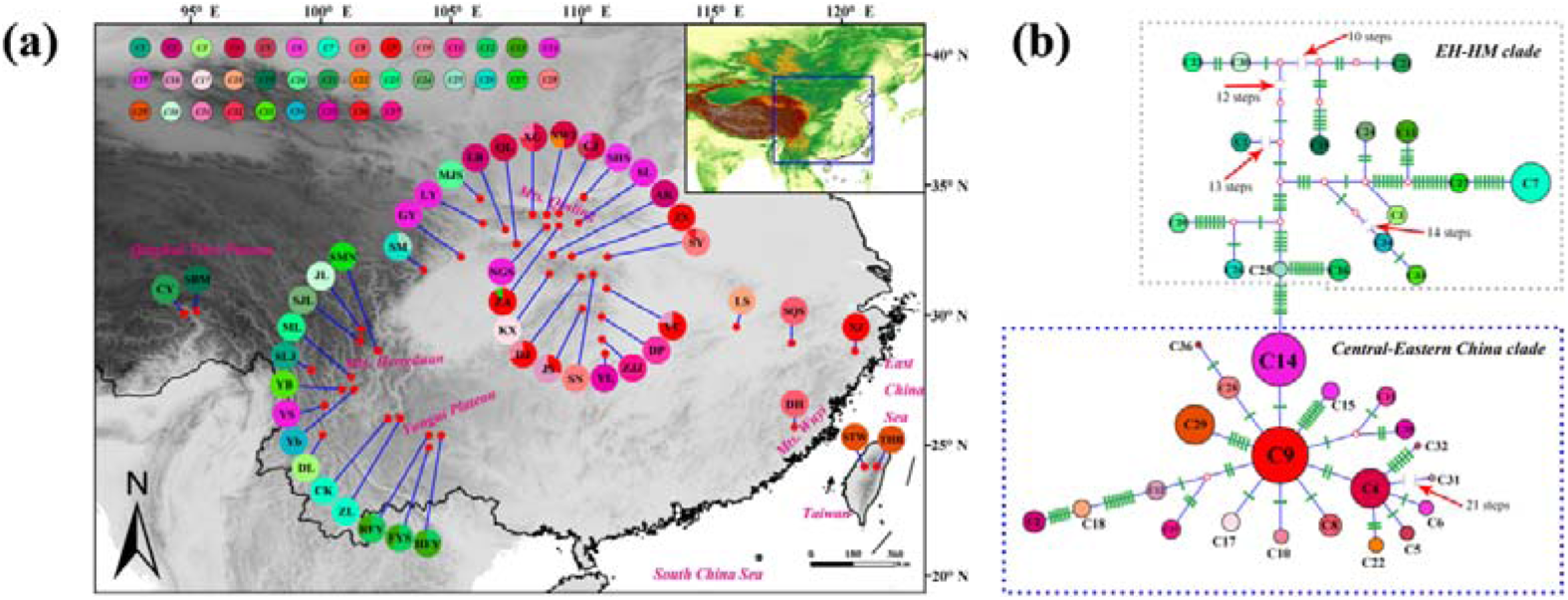
(a) Geographic distribution of the chloroplast (cp) DNA haplotypes in the 46 *Quercus spinosa* populations from subtropical China. (a) Geographic distribution of haplotypes. Haplotype frequencies for each population are denoted in the pie charts and population codes are presented in the center of the pie chart (see Table S1 for population codes). (b) Genealogical relationships between the 37 cpDNA haplotypes. Each circle sector is proportional to the frequency of each chlorotype. Small open circles represent missing haplotypes. Green bars represent nucleotide variation.

In this study, we employed an integrative approach to determine the evolutionary history and genetic divergence of *Q.sipnosa* and its response to climatic changes. The specific aims were to (i) characterize the range-wide phylogeographical patterns and genetic structure; (ii) determine the divergence times of intraspecific lineages and any underlying environmental and geographical causes; (iii) evaluate how climatic and geographical variation impact the genetic structure and divergence of *Q. spinosa*; and (iiii) reveal whether multiple refugia existed for *Q. spinosa.* We believe that knowledge of the population structure and evolutionary history of the evergreen oak species would be important to understand the complicated evolutionary history of species in subtropical China.

## Materials and methods

### Sampling and genotyping

Leaf samples were collected from 776 adult belonging to 46 natural populations of *Quercus spinosa,* covering most of its distribution range in China. All sampled individuals distanced at least 100 m from each other, and sample size varied from four to 20, depending on population size (Supporting Information Table S1). After DNA extraction, 12 nSSR loci and four cpDNA fragments were amplified (see details in Note S1).

## DNA sequence analysis

### Genetic diversity, phylogenetic analyses, and divergence time estimation

Relationships among the haplotypes obtained for *Q. spinosa* were evaluated using NETWORK v4.6 (Bandelt *et al.,* 1999). Haplotype (*He*) and nucleotide (π) diversities, and Tajima's *D* (Tajima, 1989) and Fu's (*Fs*) (Fu & Li, 1993) neutrality tests to assess possible expansions and their associated significance values were calculated in DNASP v5.00.04 (Librado & Rozas, 2009), at the population, region, and species levels. In addition, for specified clades (see results section), the average gene diversity within populations (*H_S_*), total gene diversity (*H_T_*), and the differentiation of unordered (*G*_ST_) and ordered (*N*_ST_) alleles based on 1,000 random permutations were estimated in PERMUT v1.2.1 (Pons & Petit, 1996).

Congruence among sequences of different fragments was examined with the partition homogeneity test (Farris *et al.,* 1995) as implemented in PAUP* v4.0b10 (Swofford, 2003). The HKY + G nucleotide substitution model, which was determined in JMODELTEST v1.0 (Posada, 2008), and an uncorrelated lognormal relaxed clock (Drummond *et al.,* 2005) were used to estimate the phylogenetic relationships and divergence times between lineages according to the Bayesian inference methods implemented in BEAST v1.7.5 (Drummond *et al.,* 2012). *Castanea mollissima* and *Trigonobalanus doichangensis* were used as outgroups, and a Yule process tree prior was specified. Based on fossil evidence, we set the divergence time between *T. doichangensis* and other two species (F1 in Fig. 2) at 44.8 million years ago (Ma) (±SD = 3.0 Ma), providing a 95% confidence interval (CI) of 37.2–52.3 Ma. The divergence time between *C. mollissima* and *Q. spinosa* (F2 in Fig. 2) was set 28.4 Ma (±SD = 2.2 Ma), providing a 95% CI of 23.0–33.9 Ma. The detailed calibration for each point is described in Sauquet *et al.* (2012). Three independent runs of 5 × 10^7^ Markov chain Monte Carlo (MCMC) steps were carried out, sampling at every 5,000 generations, following a burn-in of the initial 10% cycles. To confirm sampling adequacy and convergence of the chains to a stationary distribution, MCMC samples were inspected in TRACER v1.5 (http://tree.bio.ed.ac.uk/software/tracer/). Trees were visualized using FIGTREE v1.3.1 (http://tree.bio.ed.ac.uk/software/figtree/).

**Fig. 2.**
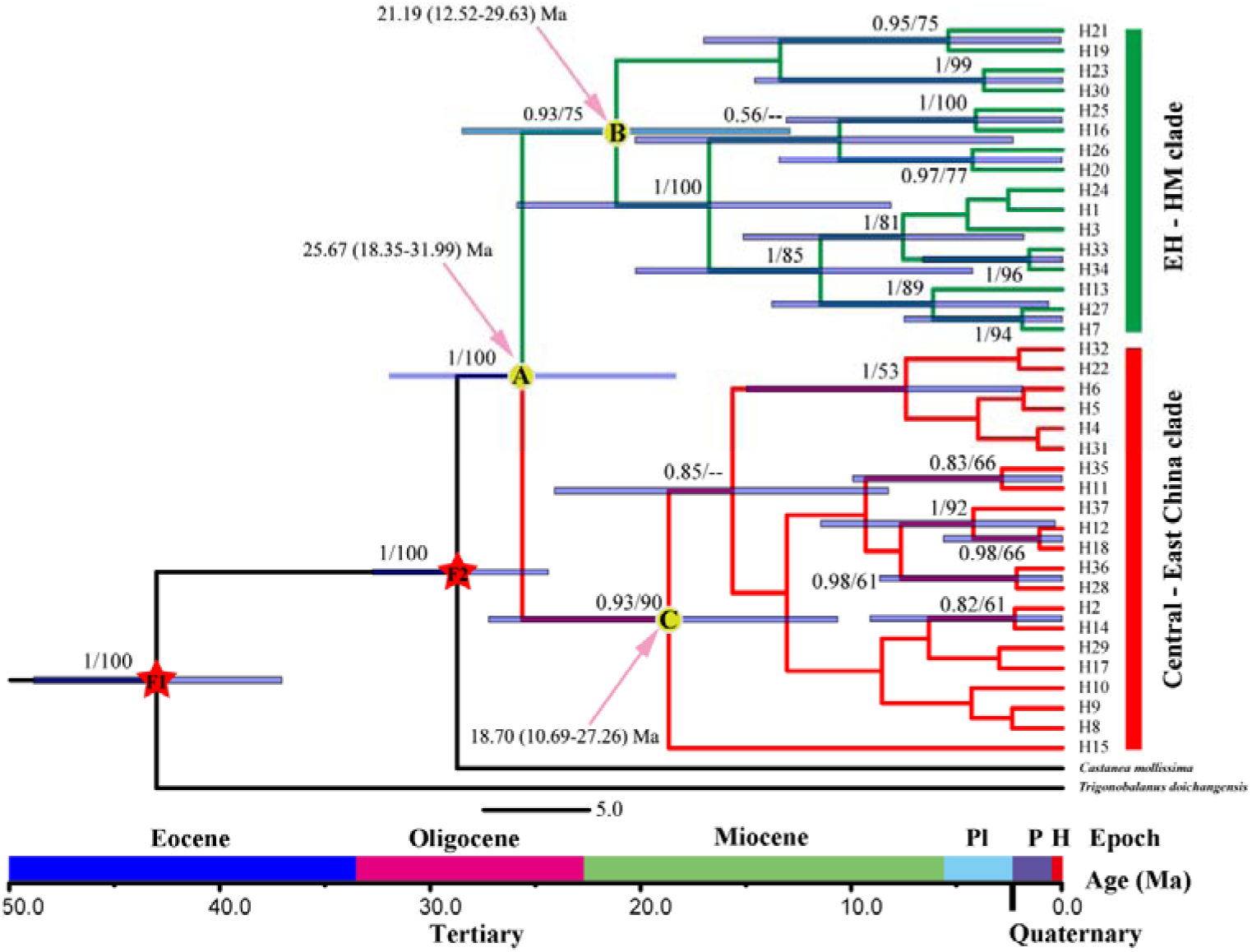
BEAST-derived chronograms of *Quercus spinosa* based on chloroplast DNA sequences (*psbA-trnH, psbB-psbF, matK,* and *Ycf1*). Red stars indicate fossil calibration points. Pink arrows indicate a recent common ancestor of *Quercus spinosa* lineages. Light blue bars indicate the 95% highest posterior density (HPD) credibility intervals for node ages (in million years ago, Ma). Bootstrap values (> 50%) based on maximum likelihood (ML) analysis and posterior probabilities are indicated above nodes.

Furthermore, haplotype phylogenetic relationships were inferred using maximum likelihood (ML), treating gaps (indels) as missing data. The ML analysis based on the HKY + G substitution model, as selected by JMODELTEST, was performed on RAXML v7.2.8 (Stamatakis *et al.,* 2008). Node support was assessed using 1,000 ‘fast bootstrap’ replicates.

### Demographic history and diversification analysis

The demographic patterns of all *Q. spinosa* populations and of each of the two groups identified in BEAST and in STRUCTURE v2.3.3 (Pritchard *et al.,* 2000) analyses (see results section) were examined through mismatch distribution analysis (MDA) in ARLEQUIN v3.5 (Excoffier & Lischer, 2010). For clades identified (see results section), we also tested the null hypothesis of spatial expansion using mismatch distribution analysis (MDA) in ARLEQUIN (see details and results in Note S2). Population demographic history was also evaluated by estimating the changes in the effective population size over time using a Bayesian skyline plot (BSP, Drummond *et al.,* 2005), as implemented in BEAST, and selecting the piecewise-linear model for tree priors. This approach incorporates uncertainty in the genealogy as it uses MCMC integration under a coalescent model. Chains were run for 100 million generations, sampling at every 10,000 generations, and their convergence and output were checked and analyzed in TRACER.

The temporal dynamics of *Q. spinosa* diversification were measured using lineages through time (LTT) plots in the APE package (Paradis *et al.,* 2004) of R v3.3. 0 (https://www.r-project.org). Plots were produced based on 100 random trees that resulted from BEAST analysis. In addition, BAMM v2.2.0 (Rabosky, 2014) was used to explore the diversification rate heterogeneity between different *Q. spinosa* groups (see results section). Analysis run for 1 × 10^7^ generations, sampling every 5,000 generations, and convergence was tested using the CODA package (Plummer *et al.,* 2006) in R, the first 10% as burn-in. Effective sample sizes were above 1,000 for all estimated parameters. The results were used to calculate diversification rates with the R package BAMMTOOLS v2.0.2 (Rabosky *et al.,* 2014).

In order to quantify genetic variations among populations and genetic clusters (as identified by STRUCTURE, NETWORK, and BEAST analyses, see below), we performed analyses of molecular variance (AMOVA) in ARLEQUIN using the *Φ-* and *R*-statistics, respectively. The significance of fixation indices was tested using 10,000 permutations (Excoffier *et al.,* 1992).

## Microsatellite data analysis

### Population genetic analysis

We used MICROCHECKER v2.2.3 (Van Oosterhout *et al.,* 2004) to test the presence of null alleles in all loci. POPGENE v1.31 (Yeh *et al.,* 1999) was used to estimate the total number of alleles (*A*_O_), observed heterozygosity (*H*_O_), expected heterozygosity over all populations (*H*_E_), gene diversity within populations (*H*_S_), and total genetic diversity (*H*_T_). Linkage disequilibrium (LD) and departure from Hardy-Weinberg equilibrium (HWE) were evaluated using FSTAT v2.9.3 (Goudet, 2001). Significance levels were corrected by the sequential Bonferroni method (Rice, 1989).

Genetic differentiation among populations was evaluated using θ (*F*_ST_) (Weir & Cockerham, 1984) and *G*’_ST_ (Hedrick, 2005) across loci in SMOGD (Crawford, 2010). Compared to traditional measures, *G*’_ST_ is a more suitable measure for highly polymorphic markers such as microsatellites.

Genetic groups were inferred using the Bayesian clustering approach implemented in STRUCTURE, based on the admixture model with independent allele frequencies. Two alternative methods were utilized to estimate the most likely number of genetic clusters (*K*) in STRUCTURE HARVESTER (Earl & vonHoldt, 2012), i.e., by tracing changes in the average of log-likelihood (L(*K*), Pritchard *et al.,* 2000) and by calculating delta *K* (*∆K,* Evanno *et al.,* 2005). Twenty independent simulations (1 ≤ *K* ≤ 20) with 5.0 × 10^5^ burn-in steps followed by 5.0 × 10^5^ MCMC steps were run. These long burn-in and run lengths, along with the large number of replicates, ensured the reproducibility of the STRUCTURE results (Gilbert *et al.,* 2012). These parameters were used in the STRUCTURE analysis conducted for each *Q. spinosa* group, except for 1 ≤ *K* ≤ 18 in the EH-HM group (see results section). The estimated admixture coefficients (Q matrix) over the 20 runs were averaged using CLUMPP v1.1 (Jakobsson & Rosenberg, 2007). Graphics were produced using DISTRUCT v1.1 (Rosenberg, 2004).

As Meirmans (2012) noted, tests of isolation by distance (IBD) may be strongly biased by the hierarchical population structure as revealed in STRUCTURE analysis. A false positive relationship between genetic distance and geographical distance among populations might be produced when population structure is not properly accounted for. In order to avoid this problem, we used a stratified Mantel test in which the locations of the populations within each putative cluster identified by STRUCTURE were permuted: 10,000 random permutations were performed between the matrix of pairwise genetic distances calculated as *F_ST_*/(1 – *F_ST_*), and that of geographic distances, using the package Vegan (Oksanen *et al.,* 2013) in R.

To estimate historical and contemporary gene flow between the clusters revealed in STRUCTURE (EH-HM and CEC clusters) and between the sub-clusters within each of this clusters, we used the programs MIGRATE-N v3.6 (Beerli, 2006) and BAYESASS v3.0 (Wilson & Rannala, 2003), respectively (see details and results in Note S2).

To predict putative barriers to gene flow, we used BARRIER v2.2 (Manni *et al.,* 2004) to find the limits associated with the highest rate of genetic change according to Monmonier’s maximum difference algorithm. We obtained 1,000 Nei’s genetic distance matrices from microsatellite data using MICROSATELLITE ANALYZER v4.05 (Dieringer & Schlötterer, 2003). In addition, the program 2MOD v0.2 (Ciofi *et al.,* 1999) was used to evaluate the relative likelihood of migration-drift equilibrium, i.e., the relative contribution of gene flow vs. genetic drift to the current population structure. After 100,000 iterations and discarding the first 10% as burn-in, the Bayes factor was obtained as *P* (gene flow) / *P* (drift).

### Tests of dynamic history by ABC modeling

Based on the STRUCTURE results for the EH-HM and the CEC clusters, five lineages were identified (see Supporting Information Fig. S1): pop1, pop2, and pop3 within the CEC cluster and pop4 and pop5 within the EH-HM cluster (see results section). As 120 different scenarios can be tested for five populations, we narrowed this number by defining nested subsets of competing scenarios that were analyzed sequentially (Table S9).

Firstly, the relationships among the three CEC populations (i.e., pop1, pop2, and pop3) were investigated. Ten possible scenarios were tested (Step 1 in Fig. 5) and the most plausible scenario (i.e., scenario 2) was chosen, then to the five populations’ analysis. In total, 15 alternative scenarios of population history were summarized for the lineages and tested using the ABC procedure (Beaumont *et al*., 2002) in DIYABC v2.0.3 (Cornuet *et al.,* 2008; 2014).

In all ABC-related analyses, uniform priors were assumed for all parameters (Table S10) and a goodness-of-fit test was used to check the priors of all parameters before implementing the simulation. Following Cavender-Bares *et al.* (2011), we assumed an average generation time of 150 years for *Q. spinosa.* To select the model that best explains the evolutionary history of this species, 10 and 5 million simulations were run for all scenarios in steps 1 and 2 of the ABC analyses, respectively. The 1% simulated data closest to the observed data was used to estimate the relative posterior probabilities of each scenario via a logistic regression approach and parameters’ posterior distributions based on the most likely scenario (Cornuet *et al.,* 2008, 2014). Each simulation was summarized by the following statistics: mean number of alleles and mean genic diversity for each lineage, *F_ST_*, mean classification index, and shared allele distance between pairs of lineages.

### Ecological niche modeling

To assess the distribution shift of *Q. spinosa* during different periods, the potential habitats present at the last interglacial (LIG, 120,000 – 140,000 years BP), the LGM (21,000 years BP), mid-Holocene (MH, 6,000 BP), and under the current climate conditions, were estimated using the maximum entropy approach (MAXENT, Elith *et al.,* 2006; Phillips & Dudík, 2008) and a genetic algorithm for rule set production (GARP, Anderson *et al.,* 2003) (see details in Note S2).

### Dissimilar climate conditions and the impact of environmental factors on genetic structure (isolation by environment)

In order to evaluate differences in present climatic conditions between EH-HM and CEC, 20 bioclimatic variables (i.e., altitude plus the 19 environmental variables) were obtained from the 46 sampling points. Also, in order to determine the contribution of present environmental conditions to the genetic structure of *Q. spinosa,* we tested the pairwise relationships between *F_ST_* and climatic distances while controlling for the geographic distance among the 46 populations, using partial Mantel tests (‘mantel.partial’ function, R Core Team, 2015) and multiple matrix regressions (MMRR script in R; Wang, 2013). Significance was tested based on 10,000 permutations (see details in Note S2).

## Results

### CpDNA diversity and population structure

Analysis of the multiple alignment of the four cpDNA regions surveyed across 397 *Q. spinosa* individuals from 46 populations (2,091 bp total length) revealed that 82 sites were variable, corresponding to 72 substitutions and 10 indels (Supporting Information Tables S3-S6). These polymorphisms defined 37 haplotypes (C1-C37), with most of the 46 surveyed populations showing a single haplotype (Fig. 1a, Table S1). At the species level, the cpDNA data revealed high haplotype diversity (*H*_T_ = 0.978) and nucleotide diversity (*π* = 0.00538) (Table S7). The parsimony network (Fig. 1b) grouped the 37 cpDNA haplotypes into two major clades (EH-HM and CEC) separated by seven mutational steps, and suggested that C9, C14, and C25 could be the ancestral haplotypes of *Q. spinosa.*

The nonhierarchical AMOVA revealed a strong population structure at the species level (*Φ*_ST_ = 0.90, *P* < 0.001). The hierarchical AMOVA revealed that 42.35% of the genetic variation was partitioned among groups (EH-HM and CEC), 49.70% among populations within groups, and 7.95% within populations (Table 1). There were no significant phylogeographic structures at the species and region levels.

**Table 1.**
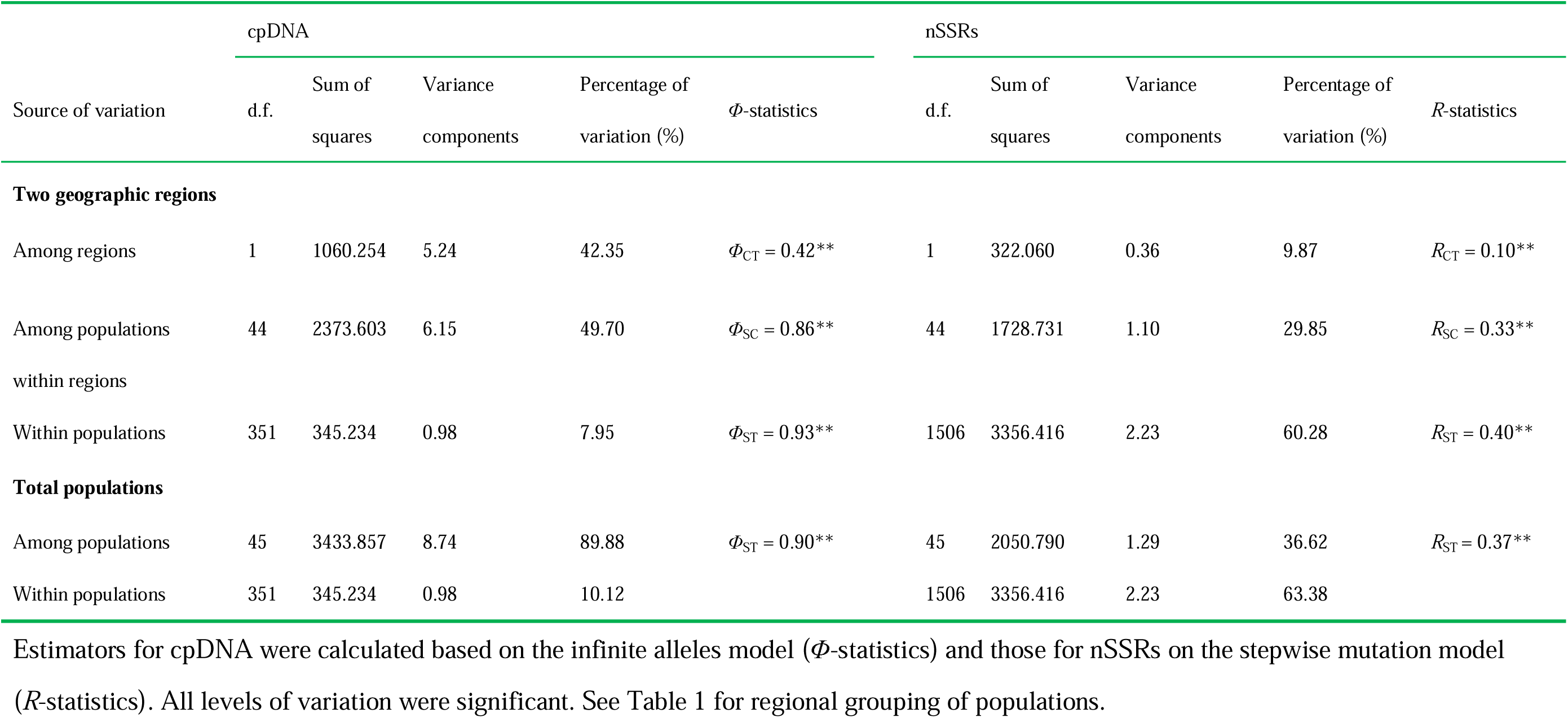
The analysis of molecular variance (AMOVA) for cpDNA data and nSSR data among two geographic regions (EH-HM and CEC) and all populations of *Quercus spinosa*

### Molecular dating, diversification rate and demography based on cpDNA data

The BEAST-derived cpDNA tree suggested *Q. spinosa* diverged from outgroup species *c.* 28.77 Ma (node F2 in Fig. 2; 95% CI: 24.43 – 32.78 Ma, PP = 1.00), indicating *Q. spinosa* and *C. mollissima* diverged during the Mid to Late Oligocene. The coalescence time estimated between the two cpDNA clades, i.e., EH-HM and CEC (25.67 Ma, node A in Fig. 2, 95% CI: 18.35 – 31.99 Ma; PP = 1.00), suggested a Late Oligocene / Early Miocene split between the two clades. Divergence times within the EH-HM (node B in Fig. 2) and CEC clades (node C in Fig. 2) were 21.19 Ma (95% CI: 12.52 – 29.63 Ma; PP = 0.93) and 18.70 Ma (95% HPD: 10.69 – 27.26 Ma; PP = 0.93), respectively. For this chronogram, BEAST provided an average substitution rate of 1.88 × 10^-10^ s/s/y, which was slower than the mean rates in other plants (e.g., 3.18 × 10^-10^ s/s/y in *Cercidiphyllum*, Qi *et al,* 2012; and 9.6 × 10^-10^ s/s/y in *Quercus glauca*, Xu *et al.,* 2014).

Our LTT analysis revealed an increase of the diversification rate of *Q. spinosa* through time (Fig. 3a). The BAMM analysis suggested a high heterogeneity in the diversification rate of the two haplotype lineages across time, with CEC presenting higher diversification rate than EH-HM (Fig. 3b).

**Fig. 3.**
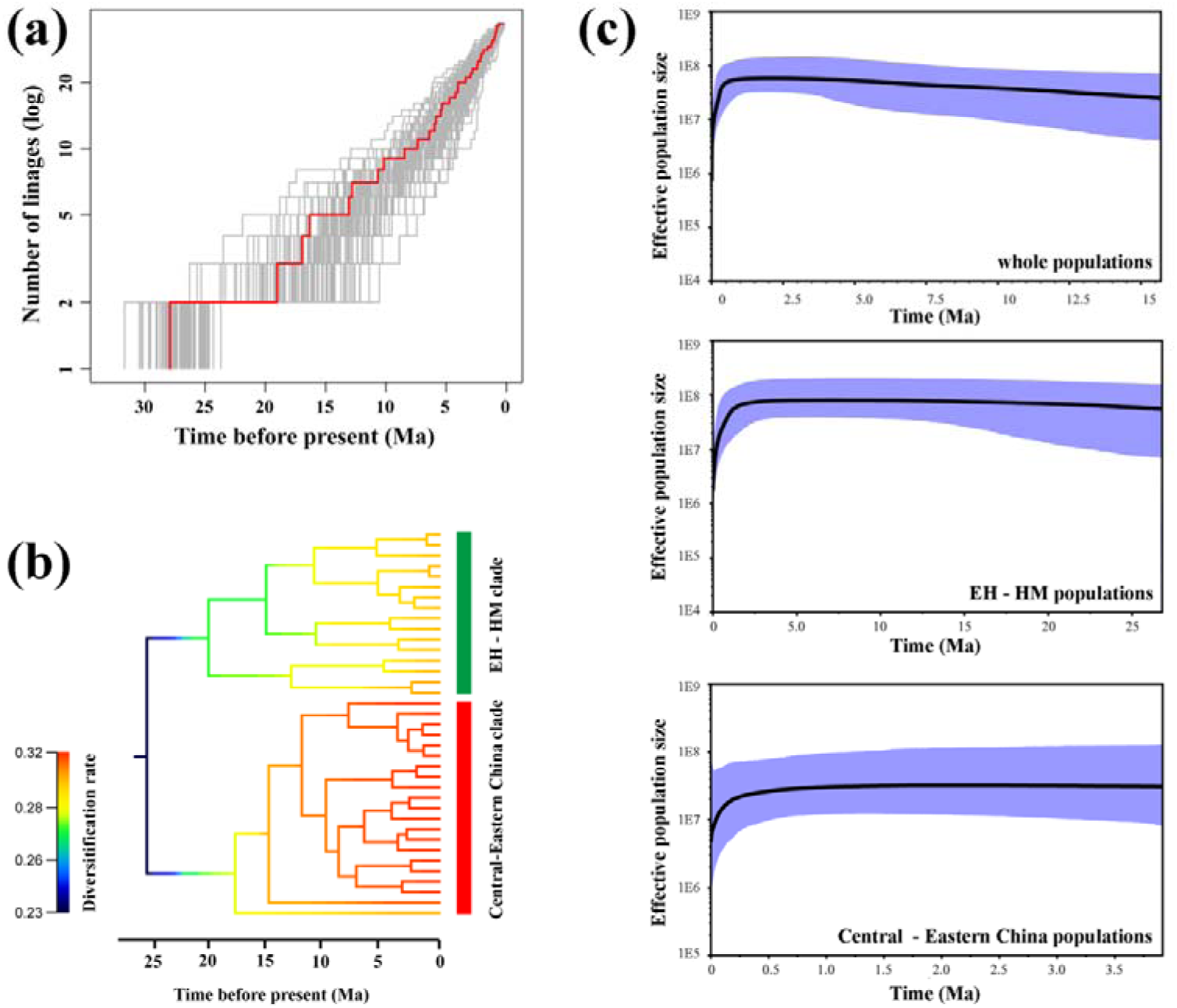
Divergence and diversification of *Quercus spinosa* through time. (a) Multiple-lineages-through-time plots based on 100 randomly sampled trees from the BEAST analysis. The red line indicates the lineages-through-time plot for the consensus chronogram. (b) Diversification rates based on chloroplast DNA haplotypes, as estimated in BAMM. (c) Bayesian skyline plot inferred from chloroplast DNA data for all *Q. spinosa* and for the two lineages, respectively. The black lines are the median effective population sizes through time and the blue areas are the limits of the 95% highest posterior densities confidence intervals.

The two clades of *Q. spinosa* generally presented non-significant Tajima’s *D* and Fu’s *F_S_* (Supporting Information Table S7). The BSP analyses indicated population sizes declined at the species and cluster levels during the Pleistocene (*c*. 0.5 Ma, 0.8 Ma, and 0.3 Ma for whole populations, EH-HM, and CEC, respectively; Fig. 3c).

### Nuclear microsatellite diversity and population structure

The null alleles test indicated a lower frequency of null alleles at each of the 12 loci than the threshold frequency (□ = 0.15) across the 46 populations, and there was no evidence for LD. After the Bonferroni corrections, significant deviation from HWE induced by homozygote excess was detected in two loci (ZAG30 and ZAG20) when all samples were treated as a single population. However, there were no HWE deviations within each population after Bonferroni correction for all nSSRs.

Screening the 776 *Q. spinosa* individuals at the 12 nSSRs revealed 160 alleles with a highly variable diversity: *A*_o_ ranged from 7 to 27, *H*_o_ from 0.091 to 0.538, *H*_S_ from 0.125 to 0.581, and *H*_T_ from 0.219 to 0.868 (Table S2). Population differentiation was significant at the 12 loci (*P* < 0.05; Table S2), with average *F*_ST_ and *G*’_ST_ reaching 0.377 and 0.573, respectively. The values of *A*_R_, *H*_o_, *H*_E_, and *F*_IS_ of each population ranged from 1.250 to 4.667, 0.137 to 0.423, 0.097 to 0.534, and −0.290 to 0.450, respectively (Table S1).

According to the STRUCTURE analysis performed for all populations (species level), *K* = 2 was optimal, although the log-likelihood of the data, log_e_ P(*K*), increased with increasing *K* (Fig. 4). Thus, this genetic structure was highly congruent with the two lineages obtained in the cpDNA analysis (EH-HM and CEC, Fig. 2). The STRUCTURE analysis performed for the CEC cluster revealed loge P(K) reached a plateau when *K* > 3 and Δ*Κ* was highest for *K* = 3 (Figs. S1 and S2). Therefore, we examined the proportional membership of each individual at *K* = 3. This showed that populations located in eastern China (DH, STW, THR, and XJ) clustered into one group (pop3), while populations located in the northern part of CEC (AK, LB, LY, NWT, QL, GY, SHS, and SY) clustered into another group (pop1); the remaining populations clustered into a third group (pop2) (Fig. S1). Two populations (SQS and LS) and a few individuals showed signs of genetic admixture (Fig. 3c, Q < 0.8) and therefore were excluded from ABC and gene flow analyses. The STRUCTURE analysis performed for the EH-HM cluster revealed *K* = 2 as optimal, according to ∆*K* (Fig. S2). For *K* = 2, populations located in the western part of EH-HM (DL, CY, JL, MJS, ML, SBM, SJL, SLJ, SM, and ZL) clustered into one group (pop4) whereas populations located in the eastern part of EH-HM (CK, FYS, HFY, RFY, SMN, Yb, and YB) clustered into another group (pop5). Population YS was excluded from ABC and gene flow analyses because it had a close genetic relationship to the CEC lineage, as revealed in the STRUCTURE analysis conducted at the species level.

**Fig. 4.**
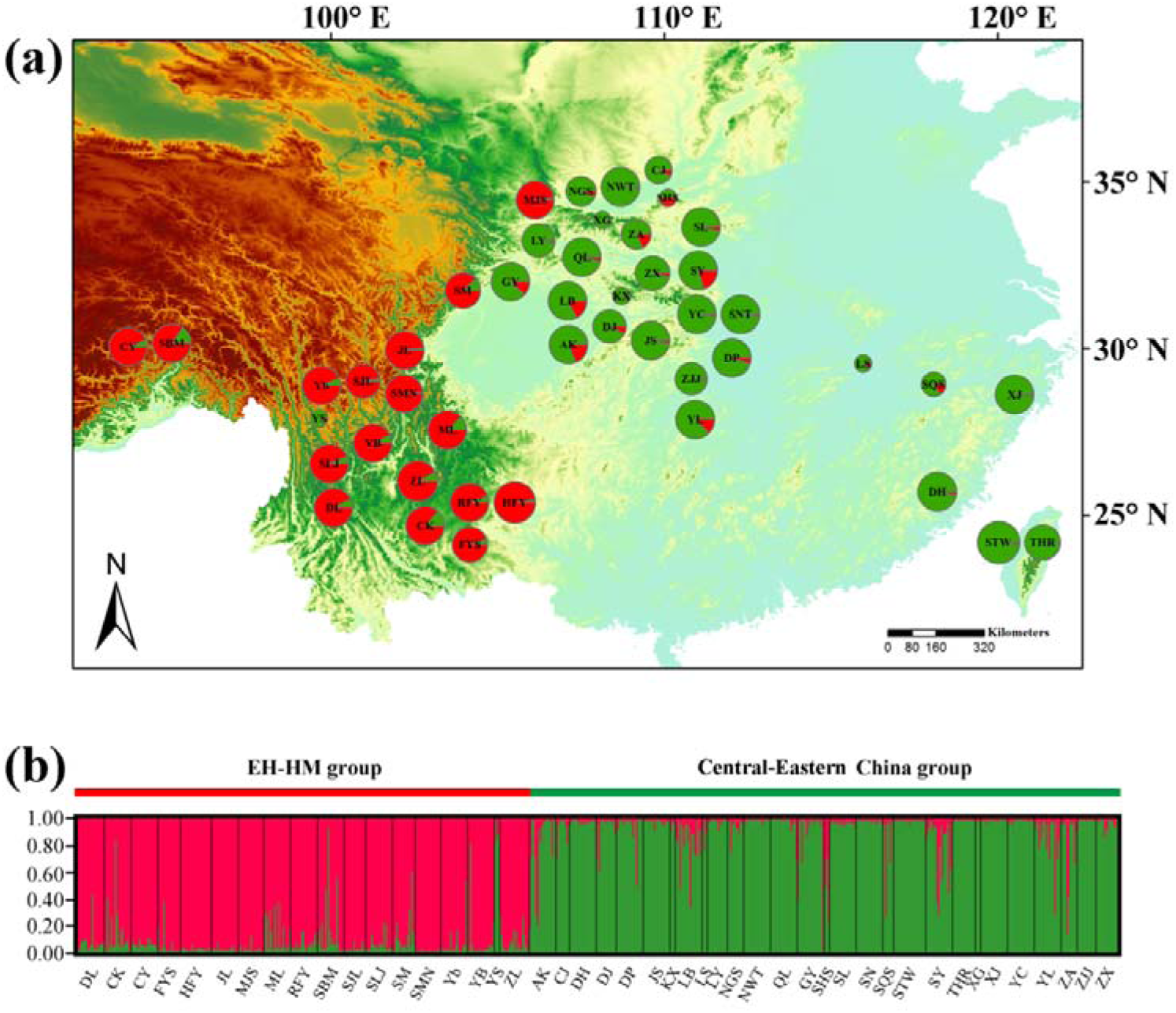
STRUCTURE analysis performed for the 46 populations of *Quercus spinosa.* (a) Geographic origin of the 46 populations and their color-coded grouping at the most likely *K* = 2. (b) Histogram of the STRUCTURE assignment test for the 46 populations based on genetic variation at 12 nuclear microsatellite loci.

The AMOVA results indicated significant genetic differentiation (*R*_ST_ = 0.40, *P* < 0.001), with only 9.87% of the variation partitioned among groups, 29.85% of the variation partitioned among populations within groups, and 68.28% of the variation partitioned among individuals within populations (Table 1). The *F*_ST_ was significant among populations within the two groups (P < 0.001), and slightly higher within CEC than within EH-HM (*F*_ST_ = 0.34 and 0.32, respectively; Table S8). There was significant IBD for *Q. spinosa* when all populations were included (r_M_ = 0.198, *P* < 0.001), and the same was found when each region was analyzed separately (r_M EH-HM_ = 0.088, *P* < 0.001; r_M_ CEC = 0.412, *P* < 0.001).

Weak genetic barriers were detected between the EH-HM and CEC lineages (bootstrap value = 14%) and within the EH-HM cluster (bootstrap values ranging from 19% to 30%) (Fig. S3). According to the 2MOD analysis, the model that most likely explains the observed population structure is the gene flow-drift model (*P* = 1.0).

### Demographic history based on nSSRs

The scenario testing results obtained in step 1 of the ABC analysis among the three eastern populations (Fig. 5) suggested scenario 2 was the most plausible, as it presented a posterior probability of 0.6913 (95% CI: 0.6399–0.7428), which was much higher than that of the other nine scenarios. In step 2, the highest posterior probability was obtained for scenario 3 (0.7564, 95% CI: 0.7102–0.8030), and this was much higher than that of the other four scenarios. According to scenario 3, the median values of the effective population sizes of pop1, pop2, pop3, pop4, pop5, and NA were 3.42 × 10^5^, 2.51 × 10^5^, 3.36 × 10^4^, 6.60 × 10^5^, 3.56 × 10^5^, and 1.19 × 10^5^, respectively. The estimated median divergence time between the EH-HM and CEC lineages (t4), within the EH-HM lineage (t3), and within the CEC lineage (t2 and t1) were 1.70 × 10^5^, 1.26 × 10^5^, 8.17 × 10^4^, and 9.22 × 10^3^ generations ago, respectively. Assuming *Q. spinosa* has a generation time of 150 years, t4, t3, t2, and t1 corresponded to 2.55 × 10^7^, 1.89 × 10^7^, 1.23 × 10^7^, and 1.38 × 10^6^ years ago, respectively. The estimated median mutation rate and proportion of multiple step mutations, based on the generalized stepwise model of microsatellites, were 2.26 × 10^-6^ and 0.485, respectively (Table 3).

**Fig. 5.**
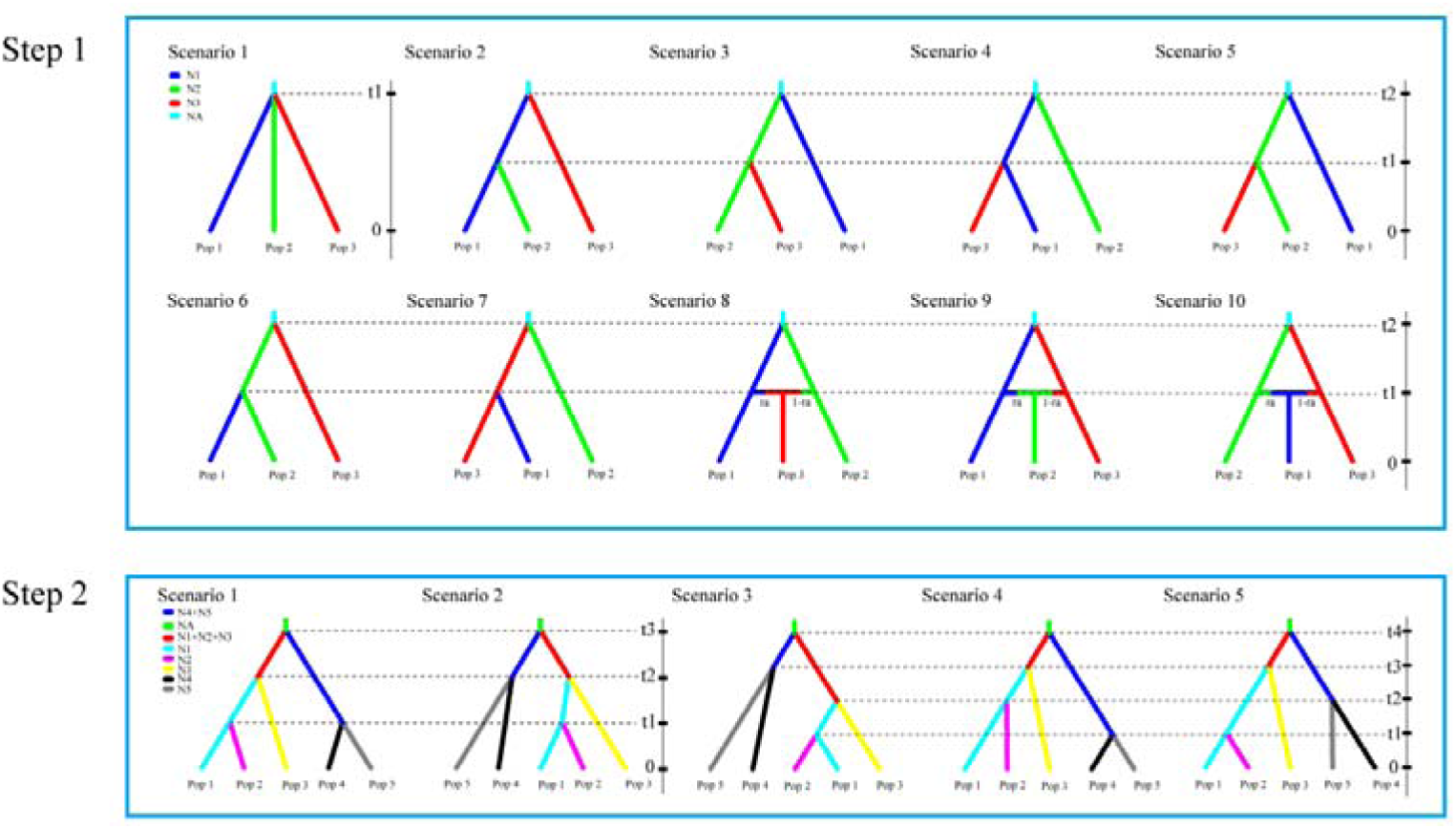
Scenarios used in the DIYABC analyses to infer the demographic history of *Quercus spinosa* based on 12 nuclear microsatellite loci. Step 1: 10 scenarios used to analyze the relationships among the three populations belonging to the Central-Eastern China lineage. Step 2: the five scenarios used to analyze the relationships among the five populations of *Q. spinosa*.

### Ecological niche modeling

There were similar change tendencies of suitable distributions of *Q.spinosa* obtained by MAXENT and GARP (Fig. 6). All models had high predictive ability (AUC > 0.9). In addition, the present-day distribution obtained for *Q. spinosa* was consistent with collection records (Fig. 6), with a potentially continuous range in the EH-HM region and western part of the CEC and a patchy distribution in eastern China. Based on our results, distribution areas during the LGM presenting moderately high suitability scores (> 0.57) significantly decreased in CCSM and MIROC compared to Holocene and present distributions, indicating a possible habitat loss during the LGM. Both procedures inferred an overall southward range shift and shrinkage of the potential distribution range during the LIG (Fig. 6e), as areas with moderately high suitability scores (> 0.57) were compressed below 30 °N and significantly decreased compared to current and MH’s distributions.

**Fig. 6.**
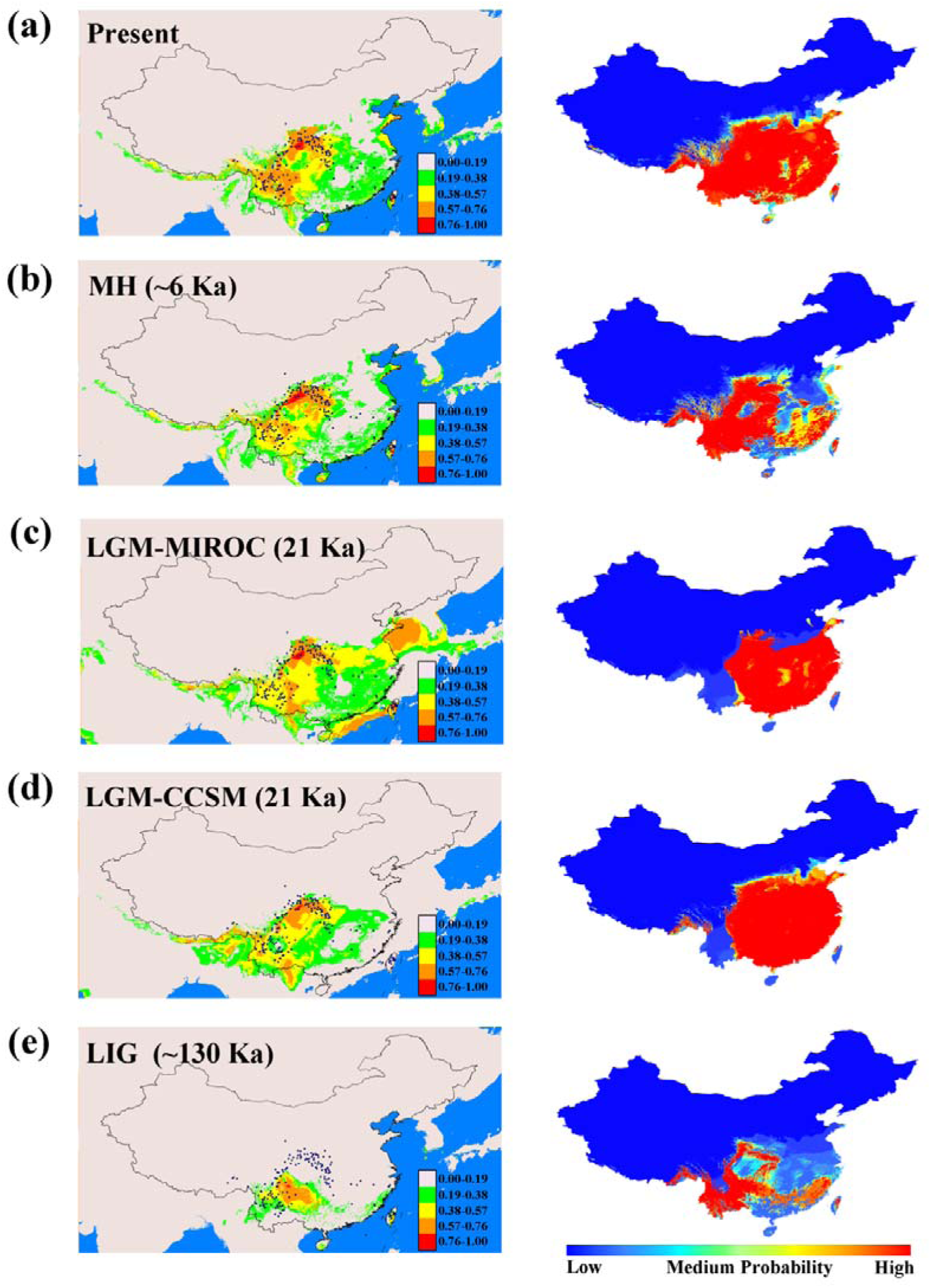
Climatically suitable areas predicted for *Quercus spinosa* using MAXENT (left) and GARP (right) in subtropical China at different times. (a) Present time; (b) Mid-Holocene (MH, *c*. 6 Ka before present (BP)); (c, d) Last glacial maximum (LGM, *c*. 21 Ka BP) under the MIROC (c) and CCSM (d) models; and (e) Last interglacial (LIG, *c*. 120–140 Ka BP). The logistic value of habitat suitability is indicated in the colored scale-bars. Black dots indicate extant occurrence points.

### Impact of the environment on *Q. spinosa* genetic structure

Climatic analyses showed that the two lineages occupied different environments, with most environmental variables significantly contributing to this divergence (Table S13 and Fig. S6). The first two PCs explained 79.21% of the variance. Whereas PC1 was mainly correlated with precipitation, PC2 was mainly correlated with temperature (Table S11). The DFA analysis suggested that 97.8% of the populations were correctly assigned to their groups (Table S12). Thus, PCA and DFA analysis clearly showed that EH-HM and CEC lineages experienced contrasting environmental conditions, presumably paving the way for divergent selection.

After controlling for geographic distance, there was a significant positive association between pairwise *F*_ST_ and PC1 at the species level and under current climatic conditions (*b*Env-PRE = 0.166, *r*Env-PRE = 0.148, *P* < 0.05); no significant relationships were obtained between pairwise *F_ST_* and PC2 (Table 2). When analyzed separately, BIO4 (*b*Env-PRE = 0.184, *P* < 0.001; *r*Env-PRE = 0.198, *P* < 0.01), BIO7 (*b*Env-PRE = 0.189, *r*Env-PRE = 0.186, *P* < 0.05), and BIO18 (*b*Env-PRE = 0.165, *r*Env-PRE = 0.158, *P* < 0.05) explained most of the genetic structure. Within the CEC lineage, a significant correlation was found between genetic differentiation and BIO4 (*b*Env-PRE = 0.244, *r*Env-PRE = 0.233, *P* < 0.05) and BIO7 (*b*Env-PRE = 0.210, *r*Env-PRE = 0.206, *P* < 0.05). However, there were no significant associations between genetic distance and PC1 or PC2 for the EH-HM and CEC lineages.

**Table 2.** Partial Mantel (PM) correlation (*r*) and multiple matrix regression (MMRR) coefficients (*b*) between genetic distance (*F*_ST_) and environmental variables for the present time.

|  | PRE |  |  |
| --- | --- | --- | --- |
|  | MMRR |  | PM |
| | $b_{Geo-PRE}$ | $b_{Env-PRE}$ | $r_{Env-PRE}$ |
| Whole range |  |  |  |
| $F_{ST}$-PC1/Geo | 0.360*** | **0.166\*** | **0.148\*** |
| $F_{ST}$-PC2/ Geo | 0.486*** | -0.074 ns | -0.078 ns |
| $F_{ST}$-BIO4/ Geo | 0.352*** | **0.198\*\*** | **0.184\*\*\*** |
| $F_{ST}$-BIO7/ Geo | 0.370*** | **0.189\*** | **0.186\*** |
| $F_{ST}$-BIO18/ Geo | 0.374*** | **0.165\*** | **0.158\*** |
| EH-HM lineage |  |  |  |
| $F_{ST}$-PC2/ Geo | 0.318* | -0.010 ns | -0.010 ns |
| $F_{ST}$-BIO4/ Geo | -0.007 ns | 0.430 ns | 0.299* |
| $F_{ST}$-BIO18/ Geo | 0.135 ns | 0.268 ns | 0.209* |
| Central-Eastern China lineage |  |  |  |
| $F_{ST}$-PC1/ Geo | 0.498* | 0.177 ns | 0.113 ns |
| $F_{ST}$-PC2/ Geo | 0.664*** | -0.051 ns | -0.067 ns |
| $F_{ST}$-BIO4/ Geo | 0.485*** | **0.244\*** | **0.233\*** |
| $F_{ST}$-BIO7/ Geo | 0.512*** | **0.210\*** | **0.206\*** |

BIO4, Temperature seasonality ( $SD \times 100$ ); BIO7, Temperature Annual Range (BIO5-BIO6); BIO18, Precipitation of warmest quarter. \*\*\*, $P < 0.001$ ; \*\*, $P < 0.01$ ; \*, $P < 0.05$ ; ns, not significant. Positive significant tests for both MMRR and PM tests are in bold.

## Discussion

### Demographic history of Q. spinosa

Our genetic data clearly evidenced two distinct lineages within *Q. spinosa* in subtropical China: one lineage was distributed in CEC region and the other in EH-HM region. Ancient events seemed to be retained in *Q. spinosa,* as suggested by the divergence times estimated for the inferred demographic processes. According to ABC simulations, the most likely demographic scenario for *Q. spinosa* involved an initially ancient isolation of two gene pools (EH-HM and CEC) followed by several divergence events within each lineage.

The intraspecific divergence of *Q. spinosa* dated back to 25.50 Ma (95% HPD: 10.83 – 41.70 Ma) or 25.67 Ma (95% HPD: 18.35 – 31.99 Ma) based on ABC simulations or BEAST-derived estimations of divergence time, respectively. The deep split between EH-HM and CEC, and within EH-HM (Table 3; Fig. 5), might have been triggered by the rapid uplift of the Himalayan – Tibetan plateau during the Oligocene (*c*. 30 Ma;Sun *et al.,* 2005; Wang *et al.,* 2012) and the early/mid Miocene (21–13 Ma; Searle, 2011). Together with the intensification of the central Asian aridity from the late Oligocene to the early Miocene (Guo *et al.,* 2002), both events lead to climatic changes, promoting the differentiation and diversification within *Q. spinosa.* Within the CEC lineage, pop1 and pop3 diverged 3.26 Ma (95% HPD: 1.16 – 10.89 Ma), coinciding with an increase in seasonality and aridity across Southeast Asia and with the intensification of Asian monsoons 3.6 Ma (An *et al.,* 2001). Such climatic changes might have contributed to the fragmentation of endemic populations and for their ultimate isolation. In addition, LTT and diversification analysis (Fig. 3a and 3b) suggested that diversification possibly started close to the Oligocene-Miocene boundary, with a rapid diversification occurring during the mid to late Miocene. Events occurring in the Late Miocene, like the rapid uplift of the Tibet plateau c. 10-7 Ma (Harrison *et al.,* 1992; Royden *et al.,* 2008) and the development of East Asian monsoons since the late Oligocene with several intensification periods during the Miocene (*c*. 15 Ma and 8 Ma; Wan *et al.,* 2007; Jacques *et al.,* 2011), might have altered habitats, enhancing the geographic isolation between and within EH-HM and CEC lineages and significantly influencing their diversification.

**Table 3.** Posterior median estimate and 95% highest posterior density interval (HPDI) for demographic parameters in scenarios 1 and 2 based on the nuclear multilocus microsatellite data for whole populations of *Quercus spinosa*

|  | Parameter | N1 | N2 | N3 | N4 | N5 | NA | t1<sup>a</sup> | t2<sup>a</sup> | t3<sup>a</sup> | t4<sup>a</sup> | μ | P |
| --- | --- | --- | --- | --- | --- | --- | --- | --- | --- | --- | --- | --- | --- |
| Scenario 3 | Median | 3.42×10<sup>5</sup> | 2.51×10<sup>5</sup> | 3.36×10<sup>4</sup> | 6.60×10<sup>5</sup> | 3.56×10<sup>5</sup> | 1.19×10<sup>5</sup> | 9.22×10<sup>3</sup> | 8.17×10<sup>4</sup> | 1.26×10<sup>5</sup> | 1.70×10<sup>5</sup> | 2.26×10<sup>-6</sup> | 0.485 |
|  | Lower_bound | 1.20×10<sup>5</sup> | 6.80×10<sup>4</sup> | 9.16×10<sup>3</sup> | 3.50×10<sup>5</sup> | 1.32×10<sup>5</sup> | 1.09×10<sup>4</sup> | 4.87×10<sup>3</sup> | 7.73×10<sup>3</sup> | 3.46×10<sup>4</sup> | 7.22×10<sup>4</sup> | 1.24×10<sup>-6</sup> | 0.166 |
|  | Upper_bound | 8.04×10<sup>5</sup> | 8.14×10<sup>5</sup> | 2.35×10<sup>5</sup> | 9.32×10<sup>5</sup> | 8.09×10<sup>5</sup> | 6.60×10<sup>5</sup> | 9.94×10<sup>3</sup> | 1.26×10<sup>5</sup> | 1.79×10<sup>5</sup> | 2.78×10<sup>5</sup> | 5.26×10<sup>-6</sup> | 0.843 |

<sup>a</sup> The unit of timing is generation.
μ: mutation rate (per generation per locus).
P represents the proportion of multiple step mutations in the generalized stepwise model, GSM.

Divergence and diversification of *Q. spinosa* appear to have occurred earlier than in other woody species in subtropical China. Whereas *Fagus* (*c*. 6.36 Ma; Zhang *et al.* (2013), *Cercidiphyllum* (*c*. 6.52 Ma;Qi *et al.* 2012), Asian white pine (*Pinus armandii*) (*c*. 7.41 Ma;Liu *et al.* 2014), and *Tetracentron sinense* (*c*. 7.41 Ma, Fig. 4; Sun *et al.* 2014) diverged during the late Pliocene. *Cyclocarya paliurus* diverged in the mid Miocene (*c*. 16.69 Ma; Kou *et al.* 2015), which is closer to the time estimated for *Q. spinosa* in the present study. Notwithstanding the differences in lineages divergence and diversification times evidenced above, pre-Quaternary climatic and/or geological events influenced the evolutionary history of Neogene taxa in subtropical China, including *Q. spinosa.*

The divergence time presented for *Q. spinosa* should be treated with caution as our molecular dating was influenced by the large variation associated with fossil calibration points, limited cpDNA variation (82 variable sites), and microsatellite data characteristic such as uncertain mutation models and homoplasy (Selkoe & Toonen, 2006). Takezaki & Nei (1996) suggested that homoplasy at microsatellite loci tended to underestimate divergence time over large time-scales. However, it does not represent a significant problem as it can be compensated using numerous loci (Estoup *et al.,* 2002). In addition, the assumption of no gene flow in DIYABC leads to the underestimation of the divergence time between species (Leaché *et al.,* 2013), although STRUCTURE analyses indicated little admixture between EH-HM and CEC lineages. Thus, the reliability of dating divergence events needs to be further studied using more loci. Nevertheless, the divergence time estimated from cpDNA and nSSRs was almost congruent, supporting our confidence that it reflects the real divergence time between EH-HM and CEC lineages. Additionally, considering the most ancient closely-related fossils to *Q. spinosa* were reported from the Miocene (Zhou, 1993), and recent phylogenetic studies established the origin of the major oak lineages by the end of the Eocene (*c*. 35 Ma) (Zhou, 1993; Hubert *et al.,* 2014; Grímsson *et al.,* 2015; Simeone *et al.,* 2016). Hence, it is plausible that the split between and within the two *Q. spinosa* lineages started in the Oligocene-Miocene boundary.

The divergence time presented for *Q. spinosa* should be treated with caution as our molecular dating was influenced by the large variation associated with fossil calibration points, limited cpDNA variation (82 variable sites), and microsatellite data characteristic such as uncertain mutation models and homoplasy (Selkoe & Toonen, 2006). Takezaki & Nei (1996) suggested that homoplasy at microsatellite loci tended to underestimate divergence time over large time-scales. However, it does not represent a significant problem as it can be compensated using numerous loci (Estoup *et al.,* 2002). In addition, the assumption of no gene flow in DIYABC leads to the underestimation of the divergence time between species (Leaché *et al.,* 2013), although STRUCTURE analyses indicated little admixture between EH-HM and CEC lineages. Thus, the reliability of dating divergence events needs to be further studied using more loci. Nevertheless, the divergence time estimated from cpDNA and nSSRs was almost congruent, supporting our confidence that it reflects the real divergence time between EH-HM and CEC lineages. Additionally, considering the most ancient closely-related fossils to *Q. spinosa* were reported from the Miocene (Zhou, 1993), and recent phylogenetic studies established the origin of the major oak lineages by the end of the Eocene (*c*. 35 Ma) (Zhou, 1993; Hubert *et al.,* 2014; Grímsson *et al.,* 2015; Simeone *et al.,* 2016). Hence, it is plausible that the split between and within the two *Q. spinosa* lineages started in the Oligocene-Miocene boundary.

Our ENMs analysis suggested *Q. spinosa* continued to expand is distribution range since the LIG, in line with the tests of spatial expansion for the two clades of *Q. spinosa* (Table S7). whereas the potential distribution areas of cold-tolerant species inhabiting subtropical China, such as spruce and yews, stabilized or decreased slightly from the LGM to present days (Li *et al.,* 2013; Liu *et al.,* 2013). In addition, given the scattered mountain ridges that characterize subtropical China, especially in the CEC region, it is likely that *Q. spinosa* remained both sparsely populated and spatially fragmented throughout the Quaternary. This distribution was in fact evidenced from the past and present modeling (Fig. 6). A higher level of fragmentation would be expected to result in low gene flow among populations, which, in turn, would lead to higher *F*_ST_. Accordingly, population fragmentation in *Q. spinosa* was much severer in the CEC than in the EH-HM lineage, the *F*_ST_ value of CEC (0.343) was slightly higher than that of EH-HM (0.321) (Table S8), and gene flow within EH-HM was significantly higher than within CEC (Fig. S5). In contrast, BSP results showed a recent decrease in the effective population size of *Q. spinosa* (Fig. 3c). Despite this disagreement with ENMs, MDA and cpDNA haplotype network revealed a recent expansion of *Q. spinosa* in the CEC region. Recent simulation studies have found that the recent population declines revealed by BSP are sensitive to the hierarchical population structure and may distort the true scenario (Grant *et al.,* 2012; Heller *et al.,* 2013). Moreover, it should be noted that our population size-change estimations were based solely on the variability of the four cpDNA fragments; more accurate estimations should be inferred using more loci (Felsenstein, 2006).

### Allopatric divergence and the impact of environmental and topographical factors on population structure

The major phylogeographic break detected in the present study based on the BEAST-derived cpDNA chronogram and on Bayesian clustering analysis is shared with other widespread temperate plants, such as *Ginkgo biloba* (Gong *et al.,* 2008), *Dysosma versipellis* (Qiu *et al.,* 2009), and *Quercus glauca* (Xu *et al.,* 2014), which clearly supported long-term isolation and allopatric divergence in subtropical China. Although the EH-HM and CEC lineages of *Q. spinosa* appear to have diverged earlier than other temperate species, its time scale was from the late Oligocene to the early Miocene, when the Tibetan plateau uplifted and central Asian aridity began to intensify. Thus, ancestral *Q. spinosa* might have been distributed in eastern and western China, allopatrically diverging in response to geographical/ecological isolation during the periods of intense climatic changes and active orogeny. A similar situation might also be responsible for the subsequent lineage divergence of pop4 and pop5 in the EH-HM region and pop1, pop2, and pop3 in the CEC region (Fig. S1). However, no significant genetic barrier was detected between the two lineages or within the CEC lineage, based on the nSSR loci (Fig. S3). The poor dispersal ability of seeds might explain the maintenance of this phylogeographic break. Most *Fagus* spp. seeds drop to the ground near parent trees and only a few may roll down on steep terrain or be dispersed by animals (e.g., jays or squirrels) over short distances (Gómez, 2003; Xiao *et al.,* 2009). This might also be the case for *Q. spinosa,* although its seed dispersal mode still needs to be studied in detail.

Recent studies have demonstrated the impacts of environmental and geographic factors on population structure (e.g., Sexton *et al.,* 2014; Wu *et al.,* 2015; Zhang *et al.,* 2016). Our analyses evidenced the significant roles of geography and climate in shaping *Q. spinosa* genetic structure (Table 2). Similar to that revealed in previous studies highlighting the importance of water availability and temperature on oak species demography (Sardans & Peñuelas, 2005; Yang *et al.,* 2009; Xu *et al.,* 2013), the present study showed the effect of precipitation (PC1) on *Q. spinosa* genetic divergence but failed to uncover the effect of temperature (PC2). However, at the lineage level, both temperature (BIO 4) and precipitation (BIO 18) influenced the divergence of CEC lineage (Table 2). However, adaptation to local environments might be biased by numerous factors (Meirmans, 2012; 2015) and it wasn’t possible to explicitly test selection based on our current data. Overall, *Q. spinosa* seems to have adapted to local environments that reinforce population genetic divergence between the two lineages, but this hypothesis requires further examination using more environment-related loci.

### Multiple refugia or refugia within refugia and long-term isolation

Previous studies suggested that many temperate plant species of subtropical China had multiple refugia during the climatic changes of Quaternary (e.g., Wang *et al.,* 2009; Shi *et al.* 2014; Wang *et al.* 2015). The major phylogeographic break found between the EH-HM and CEC regions during the pre-Quaternary, which was suggested by both cpDNA and nSSRs data analyzed in the present study, also supports the existence of multiple refugia in subtropical China, although this scenario is more plausible during interglacial than during glacial periods. The high population differentiation of cpDNA in the two lineages with most populations showing a dominant haplotype, and the subdivision of EH-HM and CEC lineages into two and three gene pools, respectively, revealed in nSSRs analysis, suggested a patterns of “refugia within refugia” for the two lineages. However, the ENMs predicted few suitable habitats for *Q. spinosa* in the CEC region during the LIG (Fig. 6). Because ENMs assume the species’ current large-scale geographical distribution is in equilibrium with the environment, as well as niche conservation over time, these models may fail to capture climatic variance and the effects of topography on microclimate (Peterson, 2003; Gavin *et al.,* 2014). Thus, ENMs might have been unable to reveal potential microrefugia for *Q. spinosa.* In addition, given the island-like genetic structure and ancient divergence between or within the two lineages revealed by cpDNA data, pointing out a long-term isolation for *Q. spinosa.* Furthermore, CEC haplotypes had a star-like distribution that was compact and with few missing haplotypes, while the EH-HM lineage had many mutational steps and sparse missing haplotypes, also indicating the long-term isolation of these two lineages. Similar results were obtained for other species occurring in subtropical China (Sun *et al.,* 2014; Xu *et al.,* 2014). Hence, *Q. spinosa* might have experienced long-term isolation among multiple refugia throughout the Quaternary, with little admixture among populations from isolated refugia in the EH-HM and CEC regions. Therefore, our results support the widely proposed hypothesis that temperate forests in subtropical China experienced long-term isolation among multiple refugia throughout the late Neogene and Quaternary (Qian & Ricklefs, 2000).

## Conclusions

The analyses of *Q. spinosa* chloroplast and nuclear DNA combined with environmental analysis and ecological niche modeling, showed that the current distribution range of this species comprises two major lineages (EH-HM and CEC) that most likely diverged through climate/tectonic-induced vicariance in the pre-Quaternary, remaining in multiple long-term refugia with little admixture during the Quaternary. Thus, pre-Quaternary environmental changes profoundly influenced the evolutionary and population demographic history of *Q. spinosa* as well as its modern genetic structure. These results support the widely accepted concept that the complex topography and climatic changes occurring in East Asia since the Neogene have provided great opportunity for allopatric divergence and speciation among temperate evergreen forest species in subtropical China. Our study also pointed out that combining phylogeography, ENMs, and bioclimatic analyses allows deep insight into the diversification and evolutionary history of species.

## Acknowledgements

[to be completed]

## Author contributions

[to be completed]

## Supporting Information

Additional supporting information may be found in the online version of this article.

**Table S1** Geographic and genetic characteristics of the 46 *Quercus spinosa* populations sampled

**Table S2** Characteristics of the 12 microsatellite loci surveyed across 46 *Quercus spinosa* populations

**Tables S3-S6** Polymorphisms detected in the four chloroplast DNA fragments

**Table S7** Results of the mismatch distribution analysis and neutrality tests based on the chloroplast DNA sequences of *Quercus spinosa*

**Table S8** Genetic diversity and genetic differentiation of the 46 *Quercus spinosa* populations

**Table S9** The 15 scenarios used for inferring the demographic history of *Quercus spinosa* in the DIY ABC analysis

**Table S10** Prior distributions for model parameters used in DIY ABC

**Table S11** Principal component analysis (PCA) of the 20 environmental variables

**Table S12** Contributions of the 20 environmental variables in the discriminant function analysis (DFA) of *Quercus spinosa*

**Table S13** ANOVA results for each of the environmental variables

**Fig. S1** Results of the genetic assignment of the EH-HM and CEC *Quercus spinosa* lineages performed on STRUCTURE

**Fig. S2** Bayesian inference of the number of clusters (*K*) of *Quercus spinosa*

**Fig. S3** BARRIER analyses and the geographic distribution of the main genetic barriers

**Fig. S4** Estimates of gene flow and migration rates

**Fig. S5** Principal component analysis (PCA) of the 20 environmental variables at present

**Fig. S6** Kernel density plots of the 20 environmental variables in the EH-HM and CEC lineages.

**Note S1** molecular methods and details of ecological niche modeling and environmental factors analysis

**Note S2** Details and results of demographic analysis for chloroplast DNA, gene flow for microsatellite data, and detail methods for ecological niche modeling and Environmental variables analysis and isolation by environment.

## References

An ZS, Kutzbach JE, Prell WL, Porter SC. 2001. Evolution of Asian monsoons and phased uplift of the Himalaya-Tibetan plateau since Late Miocene times. Nature 411: 62-66.

Anderson RP, Lew D, Peterson AT. 2003. Evaluating predictive models of species' distributions: criteria for selecting optimal models. Ecological modelling 162: 211-232.

Austerlitz F, Mariette S, Machon N, Gouyon P-H, Godelle B. 2000. Effects of colonization processes on genetic diversity: differences between annual plants and tree species. Genetics 154: 1309-1321.

Bandelt HJ, Forster P, Röhl A. 1999. Median-joining networks for inferring intraspecific phylogenies. Molecular Biology and Evolution 16: 37-48.

Beaumont MA. 2010. Approximate Bayesian computation in evolution and ecology. Annual Review of Ecology, Evolution, and Systematics 41: 379-406.

Beaumont MA, Zhang W, Balding DJ. 2002. Approximate Bayesian computation in population genetics. Genetics 162: 2025-2035.

Beerli P. 2006. Comparison of Bayesian and maximum-likelihood inference of population genetic parameters. Bioinformatics 22: 341-345.

Cavender-Bares J, Gonzalez-Rodriguez A, Pahlich A, Koehler K, Deacon N. 2011. Phylogeography and climatic niche evolution in live oaks (*Quercus series Virentes*) from the tropics to the temperate zone. Journal of Biogeography 38: 962-981.

Ciofi C, Beaumontf MA, Swingland IR, Bruford MW. 1999. Genetic divergence and units for conservation in the Komodo dragon Varanus komodoensis. Proceedings of the Royal Society of London B: Biological Sciences 266: 2269-2274.

Cornuet J-M, Pudlo P, Veyssier J, Dehne-Garcia A, Gautier M, Leblois R, Marin J-M, Estoup A. 2014. DIYABC v2.0: a software to make approximate Bayesian computation inferences about population history using single nucleotide polymorphism, DNA sequence and microsatellite data. Bioinformatics 30: 1187-1189.

Cornuet J-M, Santos F, Beaumont MA, Robert CP, Marin JM, Balding DJ, Guillemaud T, Estoup A. 2008. Inferring population history with DIY. ABC: a user-friendly approach to approximate Bayesian computation. Bioinformatics 24: 2713-2719.

Crawford NG. 2010. SMOGD: software for the measurement of genetic diversity. Molecular Ecology Resources 10: 556-557.

Denk T, Grimm Guido W. 2009. Significance of pollen characteristics for infrageneric classification and phylogeny in *Quercus* (Fagaceae). International Journal of Plant Sciences 170: 926-940.

Denk T, Grimm GW. 2010. The oaks of western Eurasia: Traditional classifications and evidence from two nuclear markers. Taxon 59: 351-366.

Dieringer D, Schlötterer C. 2003. Microsatellite analyser (MSA): a platform independent analysis tool for large microsatellite data sets. Molecular Ecology Notes 3: 167-169.

Drummond AJ, Rambaut A, Shapiro B, Pybus OG. 2005. Bayesian coalescent inference of past population dynamics from molecular sequences. Molecular Biology and Evolution 22: 1185-1192.

Drummond AJ, Suchard MA, Xie D, Rambaut A. 2012. Bayesian phylogenetics with BEAUti and the BEAST 1.7. Molecular Biology and Evolution 29: 1969-1973.

Earl D, vonHoldt B. 2012. STRUCTURE HARVESTER: a website and program for visualizing STRUCTURE output and implementing the Evanno method. Conservation Genetics Resources 4: 359-361.

Elith J, Graham CH, Anderson RP, Dudík M, Ferrier S, Guisan A, Hijmans RJ, Huettmann F, Leathwick JR, Lehmann A, et al. 2006. Novel methods improve prediction of species distributions from occurrence data. Ecography 29: 129-151.

Estoup A, Jarne P, Cornuet J-M. 2002. Homoplasy and mutation model at microsatellite loci and their consequences for population genetics analysis. Molecular Ecology 11: 1591-1604.

Evanno G, Regnaut S, Goudet J. 2005. Detecting the number of clusters of individuals using the software structure: a simulation study. Molecular Ecology 14: 2611-2620.

Excoffier L, Lischer HEL. 2010. Arlequin suite ver 3.5: a new series of programs to perform population genetics analyses under Linux and Windows. Molecular Ecology Resources 10: 564-567.

Excoffier L, Smouse PE, Quattro JM. 1992. Analysis of molecular variance inferred from metric distances among DNA haplotypes: application to human mitochondrial DNA restriction data. Genetics 131: 479-491.

Farris JS, Källersjö M, Kluge AG, Bult C. 1995. Constructing a significance test for incongruence. Systematic Biology 44: 570-572.

Felsenstein J. 2006. Accuracy of coalescent likelihood estimates: do we need more sites, more sequences, or more loci? Molecular Biology and Evolution 23: 691-700.

Fu YX, Li WH. 1993. Statistical tests of neutrality of mutations. Genetics 133: 693-709.

Gómez JM. 2003. Spatial patterns in long-distance dispersal of *Quercus ilex* acorns by jays in a heterogeneous landscape. Ecography 26: 573-584.

Gavin DG, Fitzpatrick MC, Gugger PF, Heath KD, Rodríguez-Sánchez F, Dobrowski SZ, Hampe A, Hu FS, Ashcroft MB, Bartlein PJ, et al. 2014. Climate refugia: joint inference from fossil records, species distribution models and phylogeography. New Phytologist 204: 37-54.

Gilbert KJ, Andrew RL, Bock DG, Franklin MT, Kane NC, Moore J-S, Moyers BT, Renaut S, Rennison DJ, Veen T, et al. 2012. Recommendations for utilizing and reporting population genetic analyses: the reproducibility of genetic clustering using the program structure. Molecular Ecology 21: 4925-4930.

Gong W, Chen C, Dobeš C, Fu CX, Koch MA. 2008. Phylogeography of a living fossil: Pleistocene glaciations forced *Ginkgo biloba* L.(Ginkgoaceae) into two refuge areas in China with limited subsequent postglacial expansion. Molecular Phylogenetics and Evolution 48: 1094-1105.

Goudet J 2001. FSTAT, version 2.9.3.2. A program to estimate and test gene diversities and fixation indices. [WWW document] URL http://www2.unil.ch/popgen/siftwares/fstat.htm [accessed 26 March 2007].

Grímsson F, Zetter R, Grimm GW, Pedersen GK, Pedersen AK, Denk T. 2015. Fagaceae pollen from the early Cenozoic of West Greenland: revisiting Engler’s and Chaney’s Arcto-Tertiary hypotheses. Plant Systematics and Evolution 301: 809-832.

Grant WS, Liu M, Gao T, Yanagimoto T. 2012. Limits of Bayesian skyline plot analysis of mtDNA sequences to infer historical demographies in Pacific herring (and other species). Molecular Phylogenetics and Evolution 65: 203-212.

Guo ZT, Ruddiman WF, Hao QZ, Wu HB, Qiao YS, Zhu RX, Peng SZ, Wei JJ, Yuan BY, Liu TS. 2002. Onset of Asian desertification by 22 Myr ago inferred from loess deposits in China. Nature 416: 159-163.

Harrison SP, Yu G, Takahara H, Prentice IC. 2001. Palaeovegetation (Communications arising): Diversity of temperate plants in east Asia. Nature 413: 129-130.

Harrison TM, Copeland P, Kidd W, Yin A. 1992. Raising tibet. Science 255: 1663-1670.

Hedrick PW. 2005. A standardized genetic differentiation measure. Evolution 59: 1633-1638.

Heller R, Chikhi L, Siegismund HR. 2013. The confounding effect of population structure on Bayesian skyline plot inferences of demographic history. Plos One 8: e62992.

Hewitt G. 2000. The genetic legacy of the Quaternary ice ages. Nature 405: 907-913.

Hewitt GM. 2004. Genetic consequences of climatic oscillations in the Quaternary. Philosophical Transactions of the Royal Society of London. Series B: Biological Sciences 359: 183-195.

Hubert F, Grimm GW, Jousselin E, Berry V, Franc A, Kremer A. 2014. Multiple nuclear genes stabilize the phylogenetic backbone of the genus *Quercus*. Systematics and Biodiversity 12: 405-423.

Jacques FM, Guo SX, Su T, Xing YW, Huang YJ, Liu YSC, Ferguson DK, Zhou ZK. 2011. Quantitative reconstruction of the Late Miocene monsoon climates of southwest China: a case study of the Lincang flora from Yunnan Province. Palaeogeography, Palaeoclimatology, Palaeoecology 304: 318-327.

Jakobsson M, Rosenberg NA. 2007. CLUMPP: a cluster matching and permutation program for dealing with label switching and multimodality in analysis of population structure. Bioinformatics 23: 1801-1806.

Kou YX, Cheng SM, Tian S, Li B, Fan DM, Chen YG, Soltis DE, Soltis PS, Zhang ZY. 2015. The antiquity of *Cyclocarya paliurus* (Juglandaceae) provides new insights into the evolution of relict plants in subtropical China since the late Early Miocene. Journal of Biogeography 43: 351-360.

Leaché AD, Harris RB, Rannala B, Yang Z. 2013. The influence of gene flow on species tree estimation: a simulation study. Systematic Biology Doi: 10.1093/sysbio/syt049.

Li L, Abbott RJ, Liu BB, Sun YS, Li LL, Zou JB, Wang X, Miehe G, Liu JQ. 2013. Pliocene intraspecific divergence and Plio-Pleistocene range expansions within *Picea likiangensis* (Lijiang spruce), a dominant forest tree of the Qinghai-Tibet Plateau. Molecular Ecology 22: 5237-5255.

Librado P, Rozas J. 2009. DnaSP v5: a software for comprehensive analysis of DNA polymorphism data. Bioinformatics 25: 1451-1452.

Liu J, Möller M, Provan J, Gao LM, Poudel RC, Li DZ. 2013. Geological and ecological factors drive cryptic speciation of yews in a biodiversity hotspot. New Phytologist 199: 1093-1108.

Liu JQ, Sun YS, Ge XJ, Gao LM, Qiu YX. 2012. Phylogeographic studies of plants in China: advances in the past and directions in the future. Journal of Systematics and Evolution 50: 267-275.

Liu L, Hao ZZ, Liu YY, Wei XX, Cun YZ, Wang XQ. 2014. Phylogeography of *Pinus armandii* and its relatives: heterogeneous contributions of geography and climate changes to the genetic differentiation and diversification of Chinese white pines. Plos One 9: e85920.

Manni F, Guerard E, Heyer E. 2004. Geographic patterns of (genetic, morphologic, linguistic) variation: how barriers can be detected by using Monmonier's algorithm. Human biology 76: 173-190.

Mayol M, Riba M, González-Martínez SC, Bagnoli F, de Beaulieu J-L, Berganzo E, Burgarella C, Dubreuil M, Krajmerová D, Paule L, et al. 2015. Adapting through glacial cycles: insights from a long-lived tree (Taxus baccata). New Phytologist 208: 973-986.

Meirmans PG. 2012. The trouble with isolation by distance. Molecular Ecology 21: 2839-2846.

Meirmans PG. 2015. Seven common mistakes in population genetics and how to avoid them. Molecular Ecology 24: 3223-3231.

Menitskii IL, Fedorov AA. 2005. Oaks of Asia. Enfield, USA: Science Publishers.

Myers N, Mittermeier RA, Mittermeier CG, da Fonseca GAB, Kent J. 2000. Biodiversity hotspots for conservation priorities. Nature 403: 853-858.

Nosil P, Vines TH, Funk DJ. 2005. Reproductive isolation caused by natural selection against immigrants from divergent habitats. Evolution 59: 705-719.

Oksanen J, Blanchet FG, Kindt R, Legendre P, Minchin PR, O’Hara R, Simpson GL, Solymos P, Stevens MHH, Wagner H. 2013. Package ‘vegan’. Community ecology package, version 2.9.

Ortego J, Gugger PF, Sork VL. 2015. Climatically stable landscapes predict patterns of genetic structure and admixture in the Californian canyon live oak. Journal of Biogeography 42: 328-338.

Paradis E, Claude J, Strimmer K. 2004. APE: analyses of phylogenetics and evolution in R language. Bioinformatics 20: 289-290.

Peterson AT. 2003. Predicting the geography of species’ invasions via ecological niche modeling. The Quarterly Review of Biology 78: 419-433.

Petit RJ, Carlson J, Curtu AL, Loustau M-L, Plomion C, González-Rodríguez A, Sork V, Ducousso A. 2013. Fagaceae trees as models to integrate ecology, evolution and genomics. New Phytologist 197: 369-371.

Phillips SJ, Dudík M. 2008. Modeling of species distributions with Maxent: new extensions and a comprehensive evaluation. Ecography 31: 161-175.

Plummer M, Best N, Cowles K, Vines K. 2006. CODA: Convergence diagnosis and output analysis for MCMC. R news 6: 7-11.

Pons O, Petit R. 1996. Measwring and testing genetic differentiation with ordered *versus* unordered alleles. Genetics 144: 1237-1245.

Posada D. 2008. jModelTest: phylogenetic model averaging. Molecular Biology and Evolution 25: 1253-1256.

Pritchard JK, Stephens M, Donnelly P. 2000. Inference of population structure using multilocus genotype data. Genetics 155: 945-959.

Qi XS, Chen C, Comes HP, Sakaguchi S, Liu YH, Tanaka N, Sakio H, Qiu YX. 2012. Molecular data and ecological niche modelling reveal a highly dynamic evolutionary history of the East Asian Tertiary relict *Cercidiphyllum* (Cercidiphyllaceae). New Phytologist 196: 617-630.

Qian H, Ricklefs RE. 2000. Large-scale processes and the Asian bias in species diversity of temperate plants. Nature 407: 180-182.

Qiu YX, Guan BC, Fu CX, Comes HP. 2009. Did glacials and/or interglacials promote allopatric incipient speciation in East Asian temperate plants? Phylogeographic and coalescent analyses on refugial isolation and divergence in *Dysosma versipellis*. Molecular Phylogenetics and Evolution 51: 281-293.

Qiu YX, Fu CX, Comes HP. 2011. Plant molecular phylogeography in China and adjacent regions: Tracing the genetic imprints of Quaternary climate and environmental change in the world's most diverse temperate flora. Molecular Phylogenetics and Evolution 59: 225-244.

Rabosky DL. 2014. Automatic detection of key innovations, rate shifts, and diversity-dependence on phylogenetic trees. Plos One 9: e89543.

Rabosky DL, Grundler M, Anderson C, Title P, Shi JJ, Brown JW, Huang H, Larson JG. 2014. BAMMtools: an R package for the analysis of evolutionary dynamics on phylogenetic trees. Methods in Ecology and Evolution 5: 701-707.

Rainey PB, Travisano M. 1998. Adaptive radiation in a heterogeneous environment. Nature 394: 69-72.

Rice WR. 1989. Analyzing tables of statistical tests. Evolution 43: 223-225.

Rosenberg NA. 2004. Distruct: a program for the graphical display of population structure. Molecular Ecology Notes 4: 137-138.

Royden LH, Burchfiel BC, van der Hilst RD. 2008. The geological evolution of the Tibetan Plateau. Science 321: 1054-1058.

Sardans J, Peñuelas J. 2005. Drought decreases soil enzyme activity in a Mediterranean *Quercus ilex* L. forest. Soil Biology and Biochemistry 37: 455-461.

Sauquet H, Ho SYW, Gandolfo MA, Jordan GJ, Wilf P, Cantrill DJ, Bayly MJ, Bromham L, Brown GK, Carpenter RJ, et al. 2012. Testing the impact of calibration on molecular divergence times using a fossil-rich group: the case of *Nothofagus* (Fagales). Systematic Biology 61: 289-313.

Searle MP. 2011. Geological evolution of the Karakoram Ranges. Italian journal of geosciences 130: 147-159.

Selkoe KA, Toonen RJ. 2006. Microsatellites for ecologists: a practical guide to using and evaluating microsatellite markers. Ecology Letters 9: 615-629.

Sexton JP, Hangartner SB, Hoffmann AA. 2014. Genetic isolation by environment or distance: which pattern of gene flow is most common? Evolution 68: 1-15.

Sexton JP, Hufford MB, C.Bateman A, Lowry DB, Meimberg H, Strauss SY, Rice KJ. 2016. Climate structures genetic variation across a species' elevation range: a test of range limits hypotheses. Molecular Ecology 25: 911-928.

Shi MM, Michalski SG, Welk E, Chen XY, Durka W. 2014. Phylogeography of a widespread Asian subtropical tree: genetic east–west differentiation and climate envelope modelling suggest multiple glacial refugia. Journal of Biogeography 41: 1710-1720.

Simeone MC, Grimm GW, Papini A, Vessella F, Cardoni S, Tordoni E, Piredda R, Franc A, Denk T. 2016. Plastome data reveal multiple geographic origins of *Quercus* Group *Ilex*. PeerJ 4: e1897.

Stamatakis A, Hoover P, Rougemont J. 2008. A rapid bootstrap algorithm for the RAxML web servers. Systematic Biology 57: 758-771.

Sun Y, Moore MJ, Yue LL, Feng T, Chu HJ, Chen ST, Ji YH, Wang HC, Li JQ. 2014. Chloroplast phylogeography of the East Asian Arcto-Tertiary relict *Tetracentron sinense* (Trochodendraceae). Journal of Biogeography 41: 1721-1732.

Sun ZM, Yang ZY, Pei JL, Ge XH, Wang XS, Yang TS, Li WM, Yuan SH. 2005. Magnetostratigraphy of Paleogene sediments from northern Qaidam Basin, China: implications for tectonic uplift and block rotation in northern Tibetan plateau. Earth and Planetary Science Letters 237: 635-646.

Swofford DL. 2003. PAUP*: phylogenetic analysis using parsimony, version 4b10. Sunderland, MA, USA: Sinauer Associates.

Tajima F. 1989. Statistical method for testing the neutral mutation hypothesis by DNA polymorphism. Genetics 123: 585-595.

Takezaki N, Nei M. 1996. Genetic distances and reconstruction of phylogenetic trees from microsatellite DNA. Genetics 144: 389-399.

Team RC. 2015. R: A language and environment for statistical computing [Internet]. Vienna, Austria: R Foundation for Statistical Computing; 2013. Document freely available on the internet at: http://www.r-project.org.

Van Oosterhout C, Hutchinson WF, Wills DP, Shipley P. 2004. MICRO-CHECKER: software for identifying and correcting genotyping errors in microsatellite data. Molecular Ecology Notes 4: 535-538.

Wan S, Li A, Clift PD, Stuut J-BW. 2007. Development of the East Asian monsoon: mineralogical and sedimentologic records in the northern South China Sea since 20 Ma. Palaeogeography, Palaeoclimatology, Palaeoecology 254: 561-582.

Wang IJ. 2013. Examining the full effects of landscape heterogeneity on spatial genetic variation: a multiple matrix regression approach for quantifying geographic and ecological isolation. Evolution 67: 3403-3411.

Wang J, Gao P, Kang M, Lowe AJ, Huang H. 2009. Refugia within refugia: the case study of a canopy tree *(Eurycorymbus cavaleriei)* in subtropical China. Journal of Biogeography 36: 2156-2164.

Wang YH, Jiang WM, Comes HP, Hu FS, Qiu YX, Fu CX. 2015. Molecular phylogeography and ecological niche modelling of a widespread herbaceous climber, *Tetrastigma hemsleyanum* (Vitaceae): insights into Plio-Pleistocene range dynamics of evergreen forest in subtropical China. New Phytologist 206: 852-867.

Wang Y, Zheng J, Zhang W, Li S, Liu X, Yang X, Liu Y. 2012. Cenozoic uplift of the Tibetan Plateau: Evidence from the tectonic-sedimentary evolution of the western Qaidam Basin. Geoscience Frontiers 3: 175-187.

Weir BS, Cockerham CC. 1984. Estimating F-statistics for the analysis of population structure. Evolution 38: 1358-1370.

Wilson GA, Rannala B. 2003. Bayesian inference of recent migration rates using multilocus genotypes. Genetics 163: 1177-1191.

Wright S. 1931. Evolution in mendelian populations. Genetics 16: 97-159.

Wu ZY, Wu S.1998. A proposal for a new floristic kingdom (realm): the E. Asiatic Kingdom, its delineation and characteristics: Floristic characteristics and diversity of East Asian plants: proceedings of the first international symposium of floristic characteristics and diversity of East Asian plants. Springer Verlag Beijing: China Higher Education Press.

Wu ZY, Raven P, Hong DY.1999. Flora of China. Cycadaceae through Fagaceae, vol. 4. Science Press, Beijing and Missouri Botanical Garden Press, St. Louis.

Wu ZG, Yu D, Wang Z, Li X, Xu XW. 2015. Great influence of geographic isolation on the genetic differentiation of *Myriophyllum spicatum* under a steep environmental gradient. Scientific Reports 5: 15618.

Xiao ZS, Gao X, Jiang MM, Zhang ZB. 2009. Behavioral adaptation of Pallas's squirrels to germination schedule and tannins in acorns. Behavioral Ecology 20: 1050-1055.

Xu J, Deng M, Jiang XL, Westwood M, Song YG, Turkington R. 2014. Phylogeography of *Quercus glauca* (Fagaceae), a dominant tree of East Asian subtropical evergreen forests, based on three chloroplast DNA interspace sequences. Tree Genetics & Genomes 11: 1-17.

Xu XT, Wang ZH, Rahbek C, Lessard J-P, Fang JY. 2013. Evolutionary history influences the effects of water–energy dynamics on oak diversity in Asia. Journal of Biogeography 40: 2146-2155.

Yang QS, Chen WY, Xia K, Zhou ZK. 2009. Climatic envelope of evergreen sclerophyllous oaks and their present distribution in the eastern Himalaya and Hengduan Mountains. Journal of Systematics and Evolution 47: 183-190.

Yeh FC, Yang RC, Boyle TBJ. 1999. POPGENE. version 1.31. Microsoft windows-based free ware for population genetic analysis. [WWW document] URL. http://www.ualberta.ca/~fyeh/index/htm [accessed on 12 March 2012]

Zhang YH, Wang IJ, Comes HP, Peng H, Qiu YX. 2016. Contributions of historical and contemporary geographic and environmental factors to phylogeographic structure in a Tertiary relict species, *Emmenopterys henryi* (Rubiaceae). Scientific Reports 6: 24041.

Zhang ZY, Wu R, Wang Q, Zhang ZR, López-Pujol J, Fan DM, Li DZ. 2015. Comparative phylogeography of two sympatric beeches in subtropical China: Species-specific geographic mosaic of lineages. Ecology and Evolution 3: 4461-4472.

Zhou ZK. 1993. The fossil history of *Quercus*. Acta Botanica Yunnanica 15: 21-33.

